# High Efficiency Rare Earth Element Biomining with Systems Biology Guided Engineering of *Gluconobacter oxydans*

**DOI:** 10.1101/2023.02.09.527855

**Authors:** Alexa M. Schmitz, Brooke Pian, Sabrina Marecos, Mingming Wu, Megan Holycross, Esteban Gazel, Matthew C. Reid, Buz Barstow

## Abstract

The global demand for critical rare earth elements (REE) is rising^1^ with the increase in demand for sustainable energy technologies like wind turbines^2,3^, electric vehicles^2,3^, and high efficiency lighting^4^. Current processes for producing REE require high energy inputs and can produce disproportionate amounts of hazardous waste. Biological methods for REE production are a promising solution to this problem. In earlier work we identified the most important genetic mechanisms contributing to the REE-bioleaching capability of *Gluconobacter oxydans* B58^5^. Here we have targeted two of these mechanisms to generate a high-efficiency bio-mining strain of *G. oxydans*. Disruption of the phosphate-specific transport system through a clean deletion of *pstS* constitutively turns on the phosphate starvation response, yielding a much more acidic biolixiviant, and increasing bioleaching by up to 30%. Coupling knockout of *pstS* with the over-expression of the *mgdh* membrane-bound glucose dehydrogenase gene, results in up to 73% improvement of REE-bioleaching.

## Introduction

Widespread implementation of sustainable energy infrastructure is essential for mitigating climate change^1^. Rare earth elements (REE), including the lanthanides, yttrium, and cerium are critical ingredients in many current sustainable energy technologies, including wind turbine generators^2^, solid-state lighting^6^, high-strength lightweight alloys^7,8^, and battery anodes^9^; and future ones like high-temperature superconductors^10^.

However, the extraction of REE from ore has enormous environmental impacts^11,12^. The first step in REE refining involves mining and comminution of ore, followed by gravity and/or magnetic separation to concentrate the REE-bearing solids. The REE-concentrate is then subjected to a strong acid, typically sulfuric acid due to cost, or caustic soda, and then subjected to very high temperature and sometimes pressure as well to facilitate dissolution of metal ions^11^. These extraction steps result in disproportionate amounts of hazardous waste gas, water, and often radioactive waste (such as thorium from monazite ore)^13^.

A promising solution to the environmental impact of REE-extraction is bioleaching^14,15^. Bioleaching is already used commercially for the production of about 15% of the world’s copper supply, 5% of gold, and small amounts of other metals^16^. Most of these processes depend on autotrophic (chemolithotrophic) microorganisms that oxidize ferrous iron or sulfur for energy, which in turn solubilizes the targeted metal ions^17^. Bioleaching of REE has been demonstrated at laboratory scale from a variety of solid sources including concentrated virgin ore^18^, coal fly ash^19^, and recycled and end-of-life materials^20^. REE-bioleaching typically uses heterotrophic microbes that convert sugars (glucose and/or agricultural waste) into a biolixiviant, a cocktail of solid matrix-dissolving compounds primarily composed of organic acids^21^.

*Gluconobacter oxydans* B58 is one of the most promising microorganisms for bioleaching REE^20,22^. Reed *et al*.^20^ found that biolixiviant made by *G. oxydans* was more than seven times more effective at bioleaching REE from spent FCC catalyst than a comparable concentration of gluconic acid alone. Techno-economic analysis of REE-bioleaching of spent fluid cracking catalyst with *G. oxydans* demonstrated a small margin of profit, which is highly influenced by the cost of glucose and the efficiency of extraction^23^. Bioleaching efficiency can be improved by process factors including the pulp density of the REE source (the ratio of REE mass to biolixiviant volume); continuous *vs*. batch processing; and glucose concentration^23^. However, all of these factors also influence the process economics^24^.

Genomic engineering with synthetic biology offers a promising approach to improving the efficiency of bio-mining processes without greatly affecting the process economics^25^. Previously, we identified a comprehensive set of genes underlying the efficiency of biolixiviant production and REE-bioleaching efficiency by generation and screening of a *G. oxydans* B58 whole genome knockout collection^5^.

Our earlier work identified two systems of genes that control REE-bioleaching efficiency. First, disruptions in single genes of the phosphate signaling and transport system, including *pstS, pstC, pstA*, and *pstB*, all produced large improvements in bioleaching efficiency^5^.

Second, disruptions to genes involved in glucose oxidation to gluconic acid resulted in severe attenuation of bioleaching capabilities^5^. Disruption of the *mgdh* gene that codes for the membrane-bound glucose dehydrogenase (mGDH) produced a 99% reduction in REE-bioleaching^5^. Furthermore, disruption of *mgdh* results in a re-direction of glucose into cellular metabolism and growth^22^. Likewise, disruption of genes required for synthesis of the mGDH co-factor PQQ^26^, including the *pqqABCDE* operon and *tldD* and *tldE* genes also produce large reductions in REE-bioleaching in *G. oxydans*^5^.

The results of the *G. oxydans* B58 whole genome knockout collection screen suggested a first roadmap to improving REE-bioleaching efficiency with genetic engineering: take the brakes off acid production by removing the repression of phosphate-specific transport system signaling and increase the incomplete oxidation of glucose into gluconic acid and other downstream acid products through the over-expression of *mgdh*^5^. Previous work has demonstrated that over-expression of *mgdh* results in a several-fold increase in mGDH activity and production of organic acids^27^. Here we present the results of stable mutations driving each method, and the effect of their combination on the improvement of REE-bioleaching efficiency.

## Results

### Clean Deletion of Phosphate Signaling and Transport Improves REE-bioleaching by Up to 30.1%

The phosphate-specific transport system is a transmembrane protein complex located in the bacterial inner membrane. The transmembrane sub-units PstA and PstC bind to the periplasmic phosphate-binding protein, PstS, and to the cytoplasmic signaling protein, PstB^28^. A screen of the *G. oxydans* B58 whole genome knockout collection for media-acidification through incomplete glucose oxidation found that transposon disruptions of the *pstB, pstC*, and *pstS* genes increased acidification, and disruptions of *pstB* and *pstC* also increased REE-bioleaching^5^.

As a transposon disruption does not always fully eliminate gene function, we first engineered clean deletion strains of *G. oxydans* B58 for *pstB, pstC*, and *pstS* (Δ*pstB*, Δ*pstC*, and Δ*pstS*). These clean deletion strains were then grown to saturation, along with their corresponding disruption strains (δ*pstB*, δ*pstC*, and δ*pstS)*, and wild-type *G. oxydans* B58 (wt), and mixed with glucose to produce an acidic biolixiviant. All six disruption and deletion strains had a longer lag period than wild-type when grown from a single colony. However, after back-dilution this extended lag period disappeared for all three disruption strains, Δ*pstB*, and Δ*pstS*. All disruption and deletion strains generated a significantly more acidic biolixiviant than wild-type *G. oxydans* (**Figure 1A**). Biolixiviants produced by Δ*pstB* and Δ*pstS* were considerably lower in pH than that produced by the corresponding transposon disruption strains. In contrast, biolixiviant produced by a clean deletion of *pstC* was slightly less acidic than that of the disruption strain.

**Figure 1.**
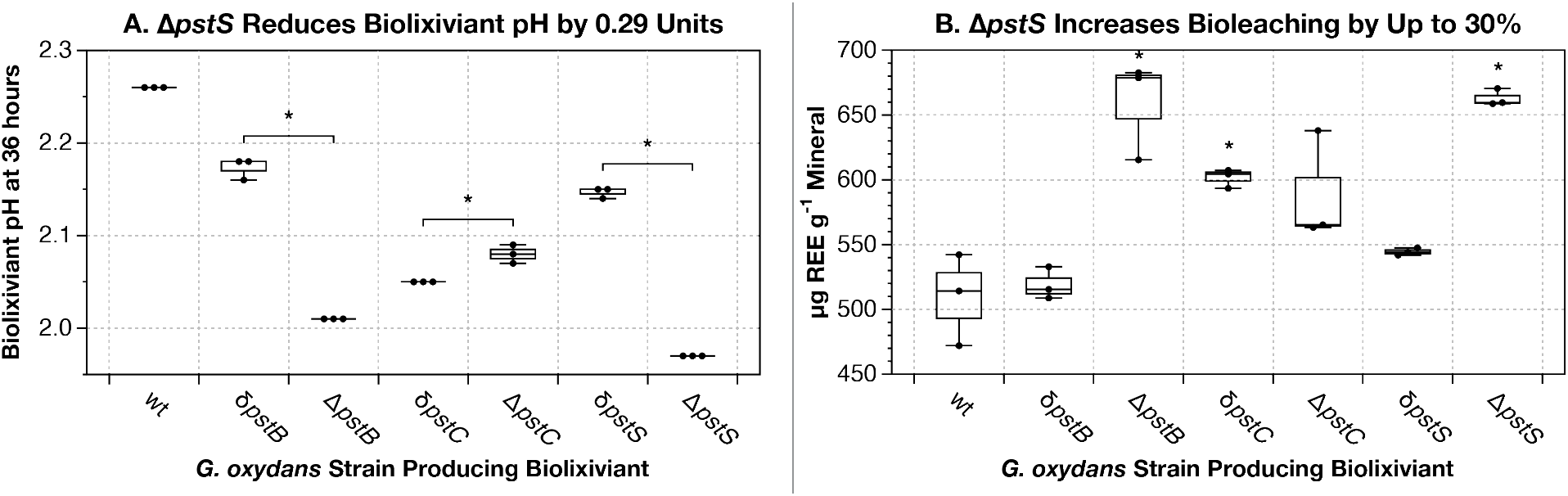
Deletion of the *pst* phosphate-specific transport genes drops biolixiviant pH by as much as 0.29 units, and improves bioleaching efficiency by up to 30.1% from allanite ore. (**A**) Effects of transposon disruption (δ) or deletion (Δ) of *pst* genes on the biolixiviant pH. Stars denote significant differences between disruption and deletion strains with a *p*-value < 0.01. The pH of the biolixiviants produced by all disruption and deletion strains were significantly different than wild type (wt) with *p*-values < 0.001. (**B**) Effects of disruption or deletion of *pst* genes on REE-bioleaching efficiency. Deletion of the ABC-type phosphate transporter ATP-binding protein PstB increases bioleaching by 29.3% over wild-type. Meanwhile, deletion of the ABC-type phosphate transporter substrate-binding protein, PstS increases bioleaching by 30.1% over wild-type. Stars denote significant improvement in total REE-bioleaching as compared with wild-type *G. oxydans, p* < 0.05. For all experiments strains were tested in triplicate, and results are demonstrative of multiple tests. Comparisons were made in Microsoft Excel with a two-tailed homoscedastic *t*-test. All data for this figure, including *p*-values, can be found in **Dataset S1**.

Biolixiviant generated by each strain was then used for REE-bioleaching from a REE-concentrated mineral ore. Biolixiviants produced by all three clean deletion strains were able to leach much more REE from the ore than wild-type (**Figure 1B**). For the disruption strains, our results were similar to previous results^5^, with the best performance coming from δ*pstC*. As expected from the higher pH of its biolixiviant, Δ*pstC* did not produce additional bioleaching improvement. Biolixiviants produced by Δ*pstB* and Δ*pstS* both greatly outperformed that of their corresponding disruption strains at REE-bioleaching. Δ*pstB* raised bioleaching by 29.3% over wild-type, while Δ*pstS* raised bioleaching by 30.1%.

### Over-expression of Membrane-bound Glucose Dehydrogenase in the Δ*pstS* Background Improves REE-bioleaching by Up to 53.1%

We hypothesized that over-expression of *mgdh* would improve both media acidification and REE-bioleaching. To test this, we selected three promoter regions previously demonstrated to confer high expression on their downstream coding regions: the *tufB* promoter^29^, and promoters P_112_ and P_114_ identified through an expression analysis of *G. oxydans* WSH-003^30,31^. Each promoter was inserted upstream of the start codon for the *mgdh* coding region to create three *mgdh* over-expression strains: P_tufB_:*mgdh*, P_112_:*mgdh*, and P_114_:*mgdh*. These insertions were also each combined with the *pstS* deletion, as it conferred the best combination of bioleaching and growth effects of the three *pst* deletion strains.

Clean deletion of *pstS* and over-expression of *mgdh* by the P_112_ promoter together had an additive effect on REE-bioleaching. In the wild-type *G. oxydans* background, P_tufB_:*mgdh* and P_114_:*mgdh* consistently produced a more acidic biolixiviant than wild-type, while P_112_:*mgdh* had no significant effect (**Figure 2A**). But, in the Δ*pstS* background, P112:*mgdh* consistently yielded the most acidic biolixiviant of all three promoter insertion strains, lowering the pH by 0.39 units. Meanwhile, P_114_:*mgdh* yielded no improvement over Δ*pstS*. Biolixiviants produced by the P_tufB_:*mgdh*, P_112_:*mgdh*, P_114_:*mgdh* all produced higher REE-bioleaching than wild-type, but none were more effective than Δ*pstS* (**Figure 2B**). When combined with the Δ*pstS* background, P_tufB_:*mgdh* and P_114_:*mgdh* were no more effective than Δ*pstS* alone. But, the combination of Δ*pstS* and P112:*mgdh* produced the most efficient REE-bioleaching strain tested (**Figure 2B**). *G. oxydans* Δ*pstS*, P_112_:*mgdh* produced REE-bioleaching that was 53.1% higher than wild-type. Taking a closer look at the leaching of each individual REE, we found that the overall composition of the leached metals did not vary between strains (**Figure 2C**).

**Figure 2.**
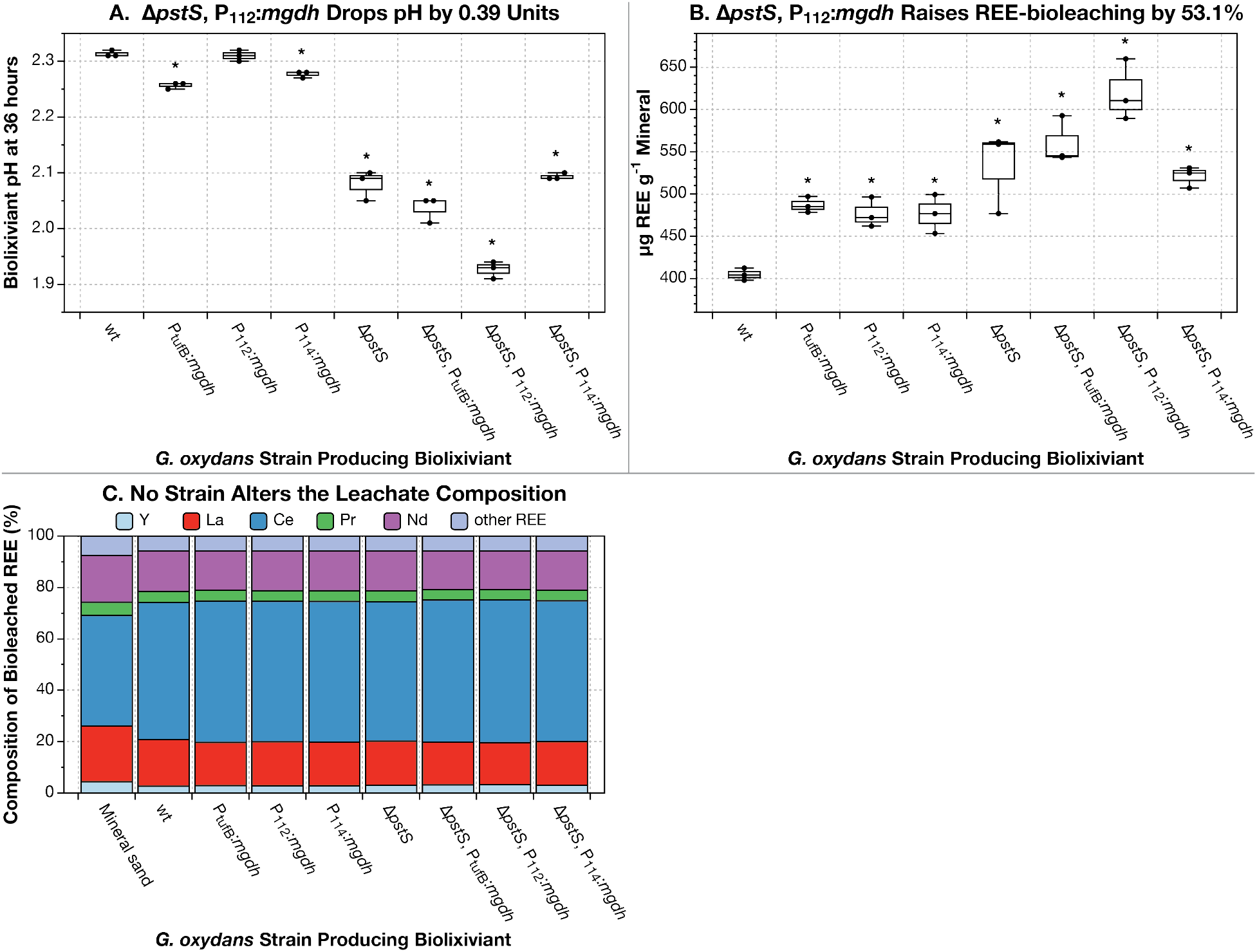
Over-expression of *mgdh* lowers biolixiviant pH by up to 0.39 units and increases REE-bioleaching from apatite sand by up to 53.1% at 10% pulp density. (**A**) Biolixiviant pH for all tested strains 36 hours after glucose introduction. (**B**) Total REE extracted per gram of apatite sand for all strains. (**C**) Total contribution of the top five REE to the total REE extracted. A pulp density of 10% was used in all bioleaching experiments (panels **B** and **C**; *i*.*e*., 10 grams of apatite in 100 mL of biolixiviant). All tests were run in triplicate and are representative of multiple experiments. Stars denote significant difference compared with wild type (wt) *G. oxydans* by a two-tailed, homoscedastic *t*-test, *p* < 0.05. All data for this figure, including *p*-values, can be found in **Dataset S2**.

### Lowering the Pulp Density to 1% Raises REE-bioleaching by G. oxydans Δ*pstS*, P_112_*:mgdh* to 73.1%

The overall efficiency of REE-bioleaching can be greatly influenced by a variety of process variables, most importantly the pulp density^23,24^. To test how these variables affect the REE-bioleaching improvement conferred by genetic engineering, we compared REE-bioleaching efficiency of the Δ*pstS* and Δ*pstS*, P_112_:*mgdh* at 1 and 10% pulp density of REE-containing mineral. At the lower pulp density, bioleaching by wild-type was only slightly higher, if at all. However, the bioleaching improvements of the engineered strains of *G. oxydans* were greater at the reduced pulp density. As in **Figure 2B**, at 10% pulp density, *G. oxydans* Δ*pstS*, P_112_:*mgdh* increased bioleaching over wild-type by 53% (**Figure 3A**). But, at 1% pulp density *G. oxydans* Δ*pstS*, P_112_:*mgdh* increased bioleaching over wild-type by 73.1% (**Figure 3A**).

**Figure 3.**
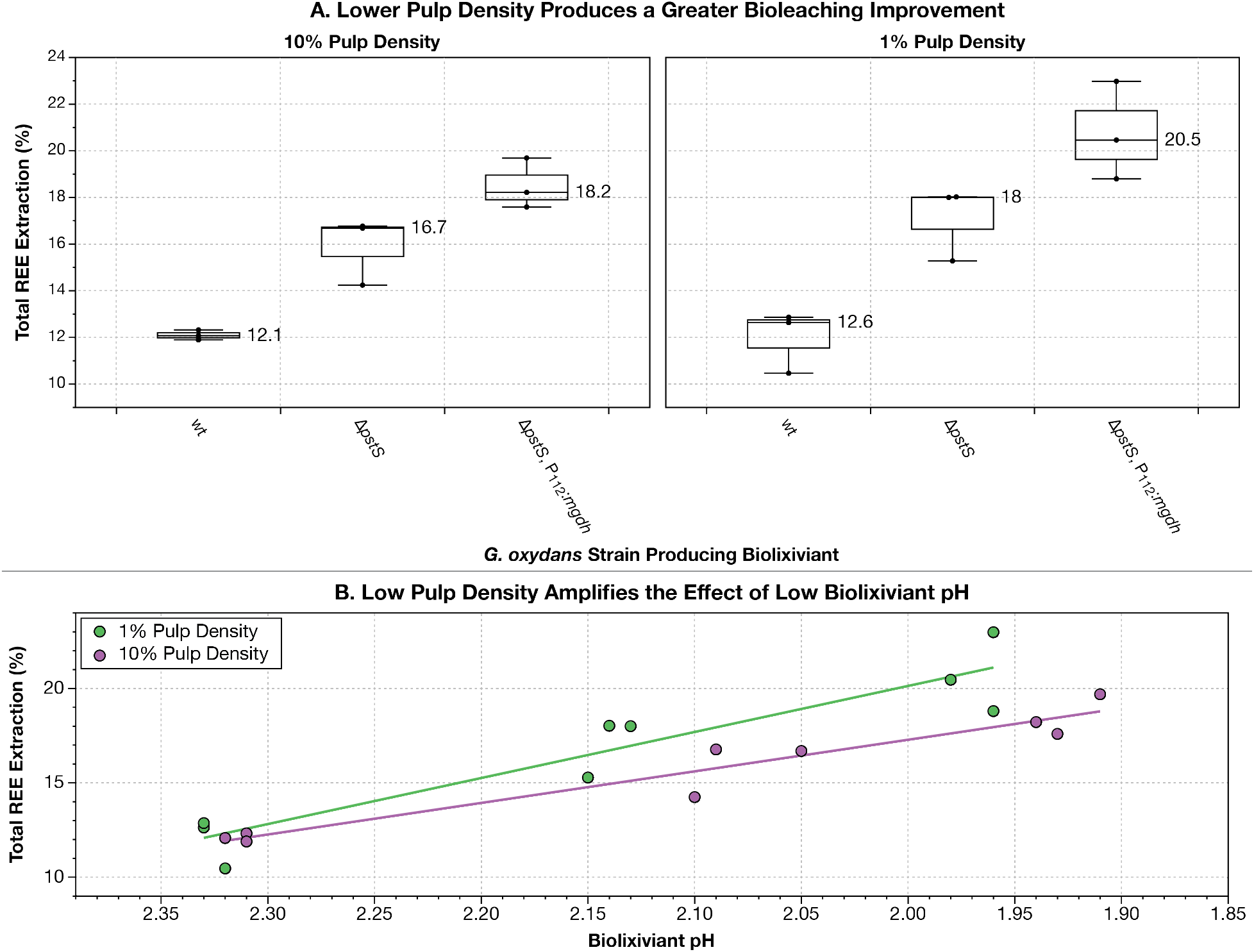
Up-regulating *mgdh* and knocking out *pstS* at the same time raises REE-extraction efficiency by up 73.1% at low pulp density. (**A**) A comparison of percent total REE-extraction at 10% (left panel) *vs*. 1% (right panel) pulp density for wild-type *G. oxydans*, Δ*pstS* and Δ*pstS*, P_112_:*mgdh*. All strains were tested in triplicate. (**B**) Correlation (green and purple lines) between biolixiviant pH and resulting percent total REE extracted at 1% (green dots) and 10% (purple dots) pulp density. All data for this figure, including *p*-values, can be found in **Dataset S3**.

Previous work with *G. oxydans* bioleaching has indicated that biolixiviant pH is a good predictor of REE-bioleaching efficiency^20,23,32^. A comparison of percent total REE extraction vs. biolixiviant pH demonstrates that the two are correlated, but that effect of lower pH is even stronger at the lower pulp density (**Figure 3B**).

## Discussion

*Gluconobacter oxydans* is an attractive candidate for the development of a high-efficiency rare earth bioleaching system. Through the incomplete oxidation of glucose, *G. oxydans* can rapidly produce a low pH biolixiviant that can be used for the solubilization of REE^20,23,32^. Additionally, the recent development of several tools for genetic engineering in *G. oxydans* has greatly increased the potential for improvement of commercially important mechanisms^30,31,33,34^. Here we have taken advantage of this genetic versatility to greatly improve REE-bioleaching through genetic engineering in *G. oxydans* B58.

The greatest single impact on REE-bioleaching came from disruption of the phosphate-specific transport system. Phosphate solubilizing microbes (PSM) such as *G. oxydans* are able to unlock inorganic phosphate from minerals in the soil through the secretion of large amounts of organic acids^35^. In *E. coli*, the deletion of *pst* genes removes repression of the *pho* regulon, resulting in a constitutive phosphate starvation response^36,37^. One of the primary mechanisms of the phosphate starvation response is the up-regulation of enzymes involved in the release and scavenging of organic phosphates^38,39^. Whether or not the *pho* regulon regulates genes in *G. oxydans* involved in the production of organic acids for mineral phosphate solubilization is still unknown, but it could explain the strong improvement of REE-bioleaching efficiency for *G. oxydans pst*-null strains.

Alternatively, a limiting factor for acid production in the wild-type bacteria may be intolerance of the increasingly acidic environment. Again in *E. coli*, the *pho* regulon has been shown to regulate genes underlying acid shock resistance, such as *asr*, which protects proteins in the periplasm from detrimental effects of low pH^40^. If the low pH of the biolixiviant is limiting to further acid production, an increase in acid shock resistance in the *pst* background would allow for great production and a lower pH biolixiviant.

To directly target up-regulation of inorganic phosphate solubilization, and thus mineral bioleaching, we inserted high-expression promoter regions directly upstream of the *mgdh* gene, which is necessary for gluconic acid production from glucose^22^. While the *tufB*, P_112_, and P_114_ promoter regions have been reported to have high expression, this may not always hold true in the increasingly acidic environment that results from biolixiviant production. We found that the three promoter regions did not perform similarly to each other, nor did they perform the same in the two different genetic backgrounds (wild-type vs. Δ*pstS*) (**Figures 2A** and **2B**).

While all three *mgdh* promoter insertions improved REE-bioleaching in the wild-type background, only P_112_:*mgdh* had a significant effect when combined with the Δ*pstS* background. A possible explanation for this difference is a differential change in promoter activity between wild-type *G. oxydans* and Δ*pstS*. Further work is needed to confirm if the increased acidification resulting from P_112_:*mgdh* in Δ*pstS* is a result of increased P_112_ promoter activity, which would indicate its regulation by *phoB*.

Our genetic edits to *G. oxydans* allow us to maximize the effect of reducing mineral pulp density on bioleaching. By applying a design of experiment model, Deng *et al*.^24^ demonstrated that pulp density is the strongest contributor to the process economics of REE-bioleaching. Using available experimental results, Deng *et al*.^24^ predicted that the optimal pulp density for maximum yearly revenue would be 50%, despite this yielding the poorest percent REE extraction. This indicated that the higher REE-bioleaching efficiency at lower pulp densities does not outweigh the added cost of producing much greater volumes of biolixiviant per unit of leached substrate.

By comparing percent REE bioleached to biolixiviant pH (**Figure 3C**), we were further able to demonstrate that driving the biolixiviant pH down can have an even stronger effect with lower pulp density. Wild-type *G. oxydans* produces the same REE-bioleaching at 1 and 10% pulp density. At 10% pulp density, the Δ*pstS*, P_112_:*mgdh* strain improved REE-bioleaching by 53% over wild-type. But, at 1% pulp density, the same strain improved bioleaching by 73%. Further modeling is needed to determine if such improvements would influence the ideal pulp density needed to maximize yearly revenue from REE-bioleaching.

## Conclusions

Through genetic engineering of targeted mechanisms underlying REE-bioleaching in *G. oxydans*, we have created a bio-mining microbe with greatly improved REE-bioleaching capability. Our highest performing strain, *G. oxydans* Δ*pstS*, P_112_:*mgdh* performs up to 73% better at REE-bioleaching than wild-type. The global demand for rare earth elements is rising with the implementation of technological innovations, especially those related to renewable energy production, storage and transmission^9^. With nearly all REE production taking place outside of the United States due to the cost of avoiding negative environmental impact^12^, the commercialization of a clean and sustainable cost-competitive process for REE extraction is in high demand. REE-bioleaching with *G. oxydans* is done at room temperature and pressure and eliminates the need for massive amounts of harmful inorganic acids.

Although further improvements are still possible, such as up-regulation of genes contributing to PQQ synthesis, the Δ*pstS*, P_112_:*mgdh* strain greatly improves on wild-type *G. oxydans* bioleaching capabilities, which are already expected to confer a margin of profit in commercial application^23,41^. Furthermore, we have demonstrated how the modification of process parameters may further capitalize on the strain’s bioleaching improvements, thus more work is needed to understand the techno-economics. Ultimately, through the creation of a high-efficiency REE-bioleaching strain, we are a step closer to the development of a clean, sustainable REE production process capable of positively impacting the world REE market without the environmental expense.

## Materials and Methods

### Genetic Engineering of G. oxydans

In all experiments *Gluconobacter oxydans* B58 (American Type Culture Collection, Manassas, VA) was cultured in yeast peptone mannitol (YPM; 5 g L^-1^ yeast extract (C7341, Hardy Diagnostics, Santa Maria, CA), 3 g L−1 peptone (211677, BD, Franklin Lakes, NJ), 25 g L^-1^ mannitol (BDH9248, VWR Chemicals, Radnor, PA)) at 30 ºC. All genetic modifications were made using the *codA*-based markerless gene deletion through homologous recombination and counter-selection with codA in the presence of *codB*^42^. For gene deletions, the 700 base pair genomic region directly upstream from the target gene’s start codon and the 700 base pair genomic region directly downstream from the target gene’s stop codon were cloned in tandem into the pKOS6b plasmid cut with XbaI using Gibson assembly^43^ (E2611, New England Biolabs, Ipswich, MA). For insertion of promoter regions, 700 base pairs upstream and downstream from the target genes were cloned into pKOS6b sandwiching the promoter region to be inserted. Primers used for all polymerase chain reactions to clone each plasmid are listed in **Dataset S4**.

pKOS6b plasmids with homologous regions were transformed into *G. oxydans* following methods described in Mostafa *et al*.^44^. Bacteria were grown from single colony to an optical density between 0.8-0.9. Cells were harvested and washed three times with a half volume of HEPES, then resuspended in 250 µL HEPES with 20% glycerol (Bluewater Chem Group, Fort Wayne, IN). Cells were flash frozen, then thawed on ice before transformation by electroporation with 2 kV on an Eppendorf Eporator. Transformed recombinant cells were recovered overnight in 1 mL EP medium (15 g L^-1^ yeast extract, 80 g L^-1^ mannitol, 2.5 g L^-1^ MgSO_4_·7H_2_O (470301-684, Ward’s Science, St. Catherines, ON, Canada), 0.5 g L^-1^ glycerol, and 1.5 g L^-1^ CaCl_2_ (0556-500G, VWR, Radnor, PA) then planted onto YPM agar supplemented with 100 µg mL^-1^ kanamycin (kan) (IB02120, VWR). Recombinants were selected and grown overnight in YPM supplemented with 100 µg mL^-1^ kan, then plated onto YPM supplemented with 60 µg mL^-1^ 5-fluorocytosine (5-FC) (TCF0321, VWR). Colonies that emerged were then transferred onto a new YPM 5-FC agar plate for clonal isolation. Colonies were isolated from several transfers and screened for recombinants with the desired mutation using colony PCR.

### Biolixiviant Production and Bioleaching

*G. oxydans* strains were grown by inoculating 2 mL Yeast-Peptone-Mannitol (YPM) media with a single colony in a culture tube and grown for 48-72 hours until the culture reached saturation. Bacterial culture was then back-diluted to an optical density of 0.05 in 10 mL YPM in a 250 mL Erlenmeyer flask, then grown for 24 hours shaking at 250 rpm. Culture was then divided into three 100 mL flasks by pipetting 3 mL into each, then 3 mL of 40% filter sterilized glucose was added to each flask resulting in a final glucose concentration of 20%. Culture and glucose were incubated for 24 hours at 30 ºC shaking at 250 rpm. Resulting biolixiviant was transferred to a 15 mL falcon tube for pH measurement, after which 5 mL was transferred back into the same flask.

Unless otherwise specified, 500 mg REE-concentrated crushed allanite ore (Zappall from WRE) was added to each flask (10% pulp density), which was then vortexed on the highest setting until all solids were wet. Flasks were then incubated at room temperature and shaking at 200 rpm for 24 hours to facilitate bioleaching. Solids were then briefly allowed to settle before 1 mL of leachate was transferred to a 2 mL micro-centrifuge tube, which was then centrifuged for 1 min at top speed in a benchtop centrifuge to pellet any remaining solids. 500 mL of leachate was filtered through a 0.45 µm AcroPrep Advance 96-well filter plate (8029, Pall Corporation, Show Low, AZ, USA) by centrifugation at 1,500 × *g* for 5 min, then diluted 100-fold into 2% trace metal grade nitric acid (JT9368, J.T. Baker, Radnor, PA). Samples were analyzed by an Agilent 7800 ICP-MS for all REE concentrations (*m*/*z*: Sc, 45; Y, 89; La, 139; Ce, 140; Pr, 141; Nd, 146; Sm, 147; Eu, 153; Gd, 157; Tb, 159; Dy, 163; Ho, 165; Er, 166; Tm, 169; Yb, 172; and Lu, 175) using a rare earth element mix standard (67349, Sigma-Aldrich, St. Louis, MO) and a rhodium in-line internal standard (SKU04736, Sigma-Aldrich, St. Louis, MO, *m*/*z* = 103). Quality control was performed by periodic measurement of standards, blanks, and repeat samples. A pWT biolixiviant sample without bioleaching was spiked with 100 ppb REE standard and analyzed for all REE concentrations as a control. ICP-MS data were analyzed using the program MassHunter, version 4.5.

## End Notes

### Data Availability

The datasets generated during and analyzed during the current study are available on Cornell eCommons.

### Code Availability

No novel code was generated in this work.

### Materials & Correspondence

Correspondence and material requests should be addressed to B.B.. Individual strains (up to ≈ 10 at a time) are available at no charge for academic researchers. We are happy to supply a duplicate of the entire *G. oxydans* knockout collection to academic researchers, but will require reimbursement for materials, supplies and labor costs. Commercial researchers should contact Cornell Technology Licensing for licensing details.

## Author Contributions

Conceptualization: A.M.S. and B.B.; Methodology: A.M.S. and B.B.; Investigation: A.M.S., B.P., S.M. and B.B; Writing - Original Draft: A.M.S. and B.B.; Writing - Review & Editing: A.M.S., M.W., M.H., E.G., M.C.R., and B.B.; Funding Acquisition: A.M.S., M.W., M.H., E.G., and B.B.; Resources: M.C.R., E.G., and B.B.; Supervision: M.W., M.H., E.G., M.C.R, and B.B.; Data Curation: A.M.S. and B.B.; Visualization: A.M.S. and B.B.; Formal Analysis: A.M.S..

## Acknowledgements

We thank M. Weems at Western Rare Earths for advice and for gift of allanite mineral sand. A.M.S. was supported by a Cornell Energy Systems Institute Postdoctoral Fellowship, and a Small Grant from the Cornell Atkinson Center for Sustainability. This work was supported by Cornell University startup funds, an Academic Venture Fund award from the Atkinson Center for Sustainability at Cornell University, a Career Award at the Scientific Interface from the Burroughs Welcome Fund to B.B., a gift from Mary Fernando-Conrad and Tony Conrad to B.B., NSF award 2228821 to B.B. and A.M.S, and by ARPA-E award DE-AR0001341 to B.B, E.G., M.E.H, and M.W..

## Bibliography

1 The Role of Critical Minerals in Clean Energy Transitions. (International Energy Agency, 2022).

2 Dent, P. C. Rare earth elements and permanent magnets. Journal of Applied Physics 111, 07A721 (2012). https://doi.org:10.1063/1.3676616

3 Metals Demand From Energy Transition May Top Current Global Supply, <https://blogs.imf.org/2021/12/08/metals-demand-from-energy-transition-may-top-current-global-supply/> (2021).

4 Schubert, E. F. & Kim, J. K. Solid-State Light Sources Getting Smart. Science 308, 1274–1278 (2005). https://doi.org:10.1126/science.1108712

5 Schmitz, A. M. et al. Generation of a Gluconobacter oxydans knockout collection for improved extraction of rare earth elements. Nature Communications 12, 6693 (2021). https://doi.org:10.1038/s41467-021-27047-4

6 Nazarov, M. & Noh, D. New Generation of Europium- and Terbium-Activated Phosphors. (Pan Stanford Publishing, 2011).

7 Norman, A. F., Prangnell, P. B. & McEwen, R. S. The solidification behaviour of dilute aluminium–scandium alloys. Acta Materialia 46, 5715--5732 (1998).

8 Adesina, O., Anzai, I. A., Avalos, J. L. & Barstow, B. Embracing Biological Solutions to the Sustainable Energy Challenge. Chem 2, 20--51 (2017). https://doi.org:10.1016/j.chempr.2016.12.009

9 Balaram, V. Rare earth elements: A review of applications, occurrence, exploration, analysis, recycling, and environmental impact. Geoscience Frontiers 10, 1285–1303 (2019). https://doi.org:10.1016/j.gsf.2018.12.005

10 Zurek, E. & Bi, T. High-temperature superconductivity in alkaline and rare earth polyhydrides at high pressure: A theoretical perspective. J Chem Phys 150, 050901 (2019). https://doi.org:10.1063/1.5079225

11 Lucas, J., Lucas, P., Le Mercier, T., Rollat, A. & Davenport, W. Rare Earths: Science, Technology, Production and Use. (Elsevier Inc., 2014).

12 Eggert, R. et al. Rare Earths: Market Disruption, Innovation, and Global Supply Chains. Annual Review of Environment and Resources 41, 199–222 (2016). https://doi.org:10.1146/annurev-environ-110615-085700

13 Lucas, J., Lucas, P., Le Mercier, T., Rollat, A. & Davenport, W. in Rare Earths 47–67 (2015).

14 Mowafy, A. M. Biological leaching of rare earth elements. World J Microbiol Biotechnol 36, 61 (2020). https://doi.org:10.1007/s11274-020-02838-x

15 Rasoulnia, P., Barthen, R. & Lakaniemi, A.-M. A critical review of bioleaching of rare earth elements: The mechanisms and effect of process parameters. Critical Reviews in Environmental Science and Technology 51, 378–427 (2020). https://doi.org:10.1080/10643389.2020.1727718

16 Johnson, D. B. Biomining—biotechnologies for extracting and recovering metals from ores and waste materials. Current Opinion in Biotechnology 30, 24–31 (2014). https://doi.org:10.1016/j.copbio.2014.04.008

17 Barrie Johnson, D. & Hallberg, K. B. Carbon, iron and sulfur metabolism in acidophilic micro-organisms. Adv Microb Physiol 54, 201–255 (2009). https://doi.org:10.1016/S0065-2911(08)00003-9

18 Brisson, V. L., Zhuang, W.-Q. & Alvarez-Cohen, L. Bioleaching of rare earth elements from monazite sand. Biotechnology and Bioengineering 113, 339–348 (2016). https://doi.org:10.1002/bit.25823

19 Park, S. & Liang, Y. Bioleaching of trace elements and rare earth elements from coal fly ash. International Journal of Coal Science & Technology 6, 74–83 (2019). https://doi.org:10.1007/s40789-019-0238-5

20 Reed, D. W., Fujita, Y., Daubaras, D. L., Jiao, Y. & Thompson, V. S. Bioleaching of rare earth elements from waste phosphors and cracking catalysts. Hydrometallurgy 166, 34–40 (2016). https://doi.org:10.1016/j.hydromet.2016.08.006

21 Dev, S. et al. Mechanisms of biological recovery of rare-earth elements from industrial and electronic wastes: A review. Chemical Engineering Journal 397 (2020). https://doi.org:10.1016/j.cej.2020.124596

22 Krajewski, V. et al. Metabolic Engineering of Gluconobacter oxydans for Improved Growth Rate and Growth Yield on Glucose by Elimination of Gluconate Formation▿. Appl. Environ. Microb. 76, 4369–4376 (2010). https://doi.org:10.1128/aem.03022-09

23 Thompson, V. S. et al. Techno-economic and Life Cycle Analysis for Bioleaching Rare-Earth Elements from Waste Materials. ACS Sustainable Chemistry & Engineering 6, 1602–1609 (2018). https://doi.org:10.1021/acssuschemeng.7b02771

24 Deng, S. et al. Applying design of experiments to evaluate economic feasibility of rare-earth element recovery. Procedia CIRP 90, 165–170 (2020). https://doi.org:10.1016/j.procir.2020.02.005

25 Capeness, M. J. & Horsfall, L. E. Synthetic biology approaches towards the recycling of metals from the environment. Biochemical Society Transactions 48, 1367–1378 (2020). https://doi.org:10.1042/bst20190837

26 Liu, D., Ke, X., Hu, Z.-C. & Zheng, Y.-G. Improvement of pyrroloquinoline quinone-dependent d-sorbitol dehydrogenase activity from Gluconobacter oxydans via expression of Vitreoscilla hemoglobin and regulation of dissolved oxygen tension for the biosynthesis of 6-(N-hydroxyethyl)-amino-6-deoxy-α-l-sorbofuranose. J Biosci Bioeng 131, 518–524 (2021). https://doi.org:10.1016/j.jbiosc.2020.12.013

27 Meyer, M., Schweiger, P. & Deppenmeier, U. Effects of membrane-bound glucose dehydrogenase overproduction on the respiratory chain of Gluconobacter oxydans. Appl Microbiol Biotechnol 97, 3457–3466 (2013). https://doi.org:10.1007/s00253-012-4265-z

28 Hsieh, Y.-J. & Wanner, B. L. Global regulation by the seven-component Pi signaling system. Curr Opin Microbiol 13, 198–203 (2010). https://doi.org:10.1016/j.mib.2010.01.014

29 Saito, Y. et al. Cloning of genes coding for L-sorbose and L-sorbosone dehydrogenases from Gluconobacter oxydans and microbial production of 2-keto-L-gulonate, a precursor of L-ascorbic acid, in a recombinant G. oxydans strain. Appl. Environ. Microb. 63, 454–460 (1997). https://doi.org:10.1128/aem.63.2.454-460.1997 PMID - 9023923

30 Chen, Y. et al. Identification of Gradient Promoters of Gluconobacter oxydans and Their Applications in the Biosynthesis of 2-Keto-L-Gulonic Acid. Frontiers Bioeng Biotechnology 9, 673844 (2021). https://doi.org:10.3389/fbioe.2021.673844

31 Qin, Z. et al. Repurposing the Endogenous Type I-E CRISPR/Cas System for Gene Repression in Gluconobacter oxydans WSH-003. ACS Synthetic Biology 10, 84–93 (2021). https://doi.org:10.1021/acssynbio.0c00456

32 Antonick, P. J. et al. Bio- and mineral acid leaching of rare earth elements from synthetic phosphogypsum. The Journal of Chemical Thermodynamics 132, 491--496 (2019). https://doi.org:10.1016/j.jct.2018.12.034

33 Fricke, P. M. et al. A tunable l-arabinose-inducible expression plasmid for the acetic acid bacterium Gluconobacter oxydans. Applied Microbiology and Biotechnology 104, 9267–9282 (2020). https://doi.org:10.1007/s00253-020-10905-4

34 da Silva, G. A. R. et al. The industrial versatility of Gluconobacter oxydans: current applications and future perspectives. World J Microbiol Biotechnol 38, 134 (2022). https://doi.org:10.1007/s11274-022-03310-8

35 Rodríguez, H. & Fraga, R. Phosphate solubilizing bacteria and their role in plant growth promotion. Biotechnol Adv 17, 319–339 (1999). https://doi.org:10.1016/s0734-9750(99)00014-2

36 Wanner, B. L. & Chang, B. D. The phoBR operon in Escherichia coli K-12. Journal of Bacteriology 169, 5569–5574 (1987). https://doi.org:10.1128/jb.169.12.5569-5574.1987 PMID- 2824439

37 Vuppada, R. K., Hansen, C. R., Strickland, K. A. P., Kelly, K. M. & McCleary, W. R. Phosphate signaling through alternate conformations of the PstSCAB phosphate transporter. BMC Microbiol 18, 8 (2018). https://doi.org:10.1186/s12866-017-1126-z

38 Santos-Beneit, F. The Pho regulon: a huge regulatory network in bacteria. Front Microbiol 6, 402 (2015). https://doi.org:10.3389/fmicb.2015.00402

39 Yuan, J., Wu, M., Lin, J. & Yang, L. Combinatorial metabolic engineering of industrial Gluconobacter oxydans DSM2343 for boosting 5-keto-D-gluconic acid accumulation. Bmc Biotechnol 16, 42 (2016). https://doi.org:10.1186/s12896-016-0272-y

40 Sužiedėlienė, E., Sužiedėlis, K., Garbenčiutė, V. & Normark, S. The Acid-InducibleasrGene in Escherichia coli: Transcriptional Control by thephoBROperon. Journal of Bacteriology 181, 2084–2093 (1999). https://doi.org:10.1128/jb.181.7.2084-2093.1999

41 Jin, H. et al. Sustainable Bioleaching of Rare Earth Elements from Industrial Waste Materials Using Agricultural Wastes. ACS Sustainable Chemistry & Engineering 7, 15311–15319 (2019). https://doi.org:10.1021/acssuschemeng.9b02584

42 Kostner, D., Peters, B., Mientus, M., Liebl, W. & Ehrenreich, A. Importance of codB for new codA-based markerless gene deletion in Gluconobacter strains. Applied Microbiology and Biotechnology 97, 8341–8349 (2013). https://doi.org:10.1007/s00253-013-5164-7

43 Gibson, D. G. et al. Enzymatic assembly of DNA molecules up to several hundred kilobases. Nat Methods 6, 343–345 (2009). https://doi.org:10.1038/nmeth.1318

44 Mostafa, H. E., Heller, K. J. & Geis, A. Cloning of Escherichia coli lacZ and lacY genes and their expression in Gluconobacter oxydans and Acetobacter liquefaciens. Appl Environ Microbiol 68, 2619–2623 (2002). https://doi.org:10.1128/AEM.68.5.2619-2623.2002

